# Behavioral and slice electrophysiological assessment of DREADD ligand, deschloroclozapine (DCZ) in rats

**DOI:** 10.1101/2021.10.25.465454

**Authors:** Todd B. Nentwig, J. Daniel Obray, Dylan T. Vaughan, L. Judson Chandler

**Author notes:** **Declarations of interest:** None. **Corresponding Author:** L. Judson Chandler, PhD, Department of Neuroscience, Medical University of South Carolina, 30 Courtenay St, Charleston, SC 29425.

## Abstract

Designer Receptors Exclusively Activated by Designer Drugs (DREADDs) have become a premier neuroscience research tool in the past decade for their utility in providing reversible manipulations of cellular activity following experimenter-controlled delivery of a DREADD-specific ligand. However, the commonly used DREADD ligand, clozapine-N-oxide (CNO), has metabolic and off-target effects that may confound experimental results and interpretations. Moreover, CNO has relatively poor affinity for DREADD receptors, which necessitates high doses for systemic administration applications. New DREADD ligands aim to reduce metabolic and potential off-target effects while maintaining strong efficacy for the designer receptors. Recently a novel DREADD ligand, deschloroclozapine (DCZ), was shown to induce chemogenetic-mediated cellular and behavioral effects in mice and monkeys without detectable side effects. While promising, further testing of DCZ across species and experimental paradigms is warranted. The goal of the present study was to examine the effectiveness of systemic DCZ for DREADD-based chemogenetic manipulations in behavioral and slice electrophysiological applications in rats. We demonstrate that a relatively low dose of DCZ (0.1 mg/kg) supports excitatory DREADD-mediated cFos induction, DREADD-mediated inhibition of a central amygdala-dependent behavior, and DREADD-mediated inhibition of neuronal activity in a slice electrophysiology preparation. In addition, we show that this dose of DCZ does not alter gross locomotor activity or induce a place preference/aversion in control rats without DREADD expression. Together, our findings support the use of systemic DCZ for DREADD-based manipulations in rats, and provide evidence that DCZ is a superior alternative to CNO.

## Introduction

Designer Receptors Exclusively Activated by Designer Drugs (DREADDs) are a chemogenetic tool widely used in neuroscience research to reversibly manipulate neuronal activity (Armbruster et al., 2007; Urban & Roth, 2015). DREADDs are G-protein coupled receptors (GPCRs) that were mutated to shift activation from native ligands to “inert” synthetic ligands that can be experimentally administered. A variety of DREADDs are available, but the most common are the Gi based, hM4Di, and the Gq based, hM3Dq, for inhibitory and excitatory neuronal modulation, respectively (Atasoy & Sternson, 2018; Roth, 2016). DREADDs are typically expressed through viral vectors delivered locally to brain tissue or via transgenic animals possessing conditional expression of DREADDs. Successful use of DREADDs have been observed in multiple species and across a broad range of applications (Burnett & Krashes, 2016; Raper et al., 2019; Upright et al., 2018).

DREADDs offer a powerful method for manipulating cellular activity, however, a notable limitation of this technology is the ligand used for DREADD activation (Goutaudier et al., 2019). The first DREADD ligand that was widely adopted was clozapine-*N*-oxide (CNO), an “inert” metabolite of the atypical antipsychotic compound clozapine. The initial characterization of DREADDs demonstrated that CNO exhibited negligible (<1 uM) affinity for an array of GPCRs, whereas, its parent compound, clozapine displayed a broad GPCR binding profile (Armbruster et al., 2007; Jendryka et al., 2019). However, CNO has since been shown to undergo reverse-metabolism to clozapine following systemic administration (Gomez et al., 2017; Manvich et al., 2018). Furthermore, several reports have observed that systemic CNO produces off-target effects, presumably due to back-conversion into clozapine, which can penetrate the blood brain barrier (Manvich et al, 2018; MacLaren 2016). Thus, there is a growing effort to develop DREADD ligands with improved potency and specificity.

A novel DREADD ligand, deschloroclozapine (DCZ) was recently developed and displayed promising results in mice and monkeys (Nagai et al., 2020). DCZ appears to have improved binding affinity for hM4Di and hM3Dq DREADDs with reduced off-target GPCR activity compared to other DREADD ligands. In mice and monkeys, systemic DCZ administration rapidly penetrated the brain with negligible metabolite accumulation. Further, DCZ effectively supported DREADD-mediated physiological and behavioral effects at relatively doses (0.001 – 0.1 mg/kg) without apparent off-target effects. Overall, DCZ exhibits greater potency and selectivity for muscarinic DREADD receptors than other available DREADD ligands, which will likely facilitate a shift towards DCZ use in the neuroscience research community. However, based upon prior issues that arose with DREADD ligands, care must be taken to rigorously test the effects of DCZ for DREADD activation in other species and applications. Therefore, the goal of this study was to assess the efficacy of systemic DCZ for DREADD activation in rats. Our data suggest that systemic DCZ supports hM3Dq- and hM4Di-mediated cellular and behavioral effects in rats and support the use of DCZ for muscarinic DREADD receptors over existing DREADD ligands.

## Materials & Methods

### Animals

Adult male Long-Evans rats (*N* = 31) were obtained from Envigo (Indianapolis, IN). Upon arrival, animals were individually housed in standard polycarbonate cages within a temperature- and humidity-controlled vivarium, which was maintained on a reverse 12:12 light-dark cycle with lights off at 09:00. Rats were provided with Teklad 2918 (Envigo) standard chow and water *ad libitum* unless otherwise indicated. All experimental procedures were conducted with the approval of the Institutional Animal Care and Use Committee at the Medical University of South Carolina and adhered to the guidelines set forth by the National Research Council’s Guide for the Care and Use of Laboratory Animals.

### Surgical procedures and viral vectors

Rats underwent stereotaxic surgery to inject either an excitatory (AAV8.hSyn.hM3Dq.mCherry; Addgene, catalog # 50474-AAV8) or inhibitory (AAV8.hSyn.hM4Di.mCherry; Addgene, catalog # 50475-AAV8) DREADD into the central amygdala (AP: 2.2-2.4; ML: 4.2-4.3; DV: 7-6-7-8 from skull surface). For the cFos experiment, rats received unilateral injections of hM3Dq for intra-subject comparison of DCZ-induced cFos in hemispheres expressing or lacking DREADDs. All other experiments involved bilateral virus injections. Viruses were injected at a rate of 1 nL/sec for a total of 450 nL using a Hamilton syringe equipped to a micropump controller (World Precisions Instruments, Micro4, UMC4). Syringes were slowly withdrawn after a 10-minute diffusion period following injection. DREADD activation or inhibition testing began at least 4 weeks after surgery to allow adequate viral-mediated protein expression.

### Dcz-mediated excitatory DREADD Induction of cFos expression

Rats were habituated to handling and intraperitoneal (i.p.) injections of 0.9% saline for 3 consecutive days prior to injection of DCZ. Deschloroclozapine (HelloBio, catalog # HB9126) was dissolved in 0.9% saline and administered i.p. at a dose of 0.1 mg/kg for all behavioral experiments. Animals were sacrificed ~90 minutes following DCZ injection via transcardial perfusion with 1X PBS and 10% formalin. Extracted brains were post-fixed in 10% formalin for 24 hours and transferred to 30% sucrose until tissue sectioning. A cryostat was used to obtain 40um tissue sections of the central amygdala, which was stored in 1X PBS + 0.1% NaN3 until immunofluorescent labeling.

Tissue sections were washed 3 times in 1X PBS for 5 minutes each, and then permeabilized in 1X PBS 0.4% Triton-X for 30 minutes. Tissue sections were then washed in blocking buffer (1X PBS + 0.2% Triton-X + 10% Normal Donkey Serum) for 1 hour, and then incubated overnight at 4°C in blocking buffer + primary antibodies (rabbit anti-cFos, 1:1000 dilution, Synaptic Systems, catalog # 226 003; chicken anti-mCherry, 1:300 dilution, LSBio, catalog # LS-C204825). Following primary incubation, tissue is washed 10 times in 1X PBS for 1 min each and then incubated for 2 hours in blocking buffer + secondary antibodies from Jackson ImmunoResearch Laboratories (donkey anti-rabbit 488, catalog # 711-545-152; donkey anti-chicken, 1:500 dilution, catalog # 703-585-155). Upon completion of secondary incubation, tissue is washed 3 times in 1X PBS for 10 minutes each and stored in 1X PBS + 0.1% NaN3. Tissue was mounted on standard, positively charged microscope slides and coverslipped using ProLong Gold anti-fade mounting media with DAPI (Invitrogen, catalog # P36931).

Mounted tissue was imaged using a Zeiss LSM 880 confocal microscope. The 405, 488, and 561 nm lasers were used to obtain z-series images of DAPI, cFos, and hM3Dq-mCherry, respectively. Image acquisition settings (ie, laser power, gain, zoom) were identical for all images to enable proper comparison of cFos induction between hemispheres expressing or lacking hM3Dq expression. Exported image files were opened in ImageJ (Schindelin et al., 2012) and the cell counter plugin was used to quantify the number of cFos+ and mCherry+ cells. Cell counts were performed by an individual blind to the experimental conditions.

### Slice electrophysiology Recordings

Brain slices containing the central amygdala (CeA) were obtained from rats at approximately PD 180. Rats were anesthetized using isoflurane and rapidly decapitated. The brain was then promptly removed and placed into room temperature N-Methyl-D-Glucamine (NMDG) based artificial cerebral spinal fluid (aCSF) consisting of (in mM): 93 NMDG, 2.5 KCl, 1.25 NaH_2_PO_4_, 30 NaHCO_3_, 25 D-glucose, 20 HEPES, 2.5 C_5_H_9_NO_3_S, 5 ascorbic acid, 10 MgCl_2_, and 0.5 CaCl_2_. Following slice preparation using a Leica VT1000 S vibratome, slices (290 μm thick) were placed into NMDG aCSF maintained at 34°C for 15 min after which slices were incubated for at least 45 min in room temperature aCSF consisting of (in mM): 92 NaCl, 2.5 KCl, 1.25 NaH_2_PO_4_, 30 NaHCO_3_, 25 D-glucose, 20 HEPES, 2.5 C_5_H_9_NO_3_S, 5 ascorbic acid, 10 MgCl_2_, and 0.5 CaCl_2_. After incubation, slices were transferred to a submerged recording chamber held at 34°C and bathed in recording aCSF consisting of (in mM): 125 NaCl, 2.5 KCl, 25 NaHCO_3_, 10 D-glucose, 5 ascorbic acid, 1.3 MgCl_2_, and 2 CaCl_2_. The above solutions had an osmolarity of 300 – 310 mOsm and were pH adjusted (pH 7.3 – 7.43) using NaOH and HCl and were continuously aerated with 5% CO_2_/95% O_2_.

Current clamp recordings were made using a Multiclamp 700B amplifier (Molecular Devices) connected to a Windows-PC running Axograph X software. All recordings were made from visually identified hM4Di.mCherry expressing cells within the CeA. Recording electrodes (2 – 4 MΩ resistance) were filled with a potassium gluconate internal solution containing the following (in mM): 125 potassium gluconate, 20 KCl, 10 HEPES, 1 EGTA, 2 MgCl_2_, 2 Na-ATP, 0.3 Tris-GTP, and 10 phosphocreatine. The internal solution was pH adjusted (pH 7.3 – 7.43) with an osmolarity of approximately 285 mOsm. For each neuron recorded, increasing current steps (+20 pA) of 1 s duration were applied beginning from −100 pA and continuing until >6 action potentials were elicited. Using the current at which >6 action potentials were evoked, further recordings were made to determine the effects of DCZ and CNO on neuronal excitability. This was done by recording baseline excitability at 1 minute intervals for 2 minutes followed by recording excitability at 1 minute intervals for 10 minutes during perfusion with increasing concentrations of CNO or DCZ. Recordings from multiple neurons were obtained and averaged for each animal.

### Chemogenetic inhibition of CeA neurons during dependence-escalated ethanol drinking

Rats underwent a well-characterized model of ethanol dependence and withdrawal to assess dependence-induced escalated ethanol intake. This model has three consecutive phases: baseline ethanol drinking, chronic intermittent ethanol (CIE) exposure, and ethanol drinking during withdrawal. During baseline ethanol drinking, rats are given 24 hr access to voluntarily consume 20% ethanol and water in their homecage for 4 weeks under an intermittent access 2-bottle choice (2BC) schedule (Simms et al., 2008). Rat then underwent CIE exposure, which involved 14 consecutive days of ethanol exposure by intermittent vapor inhalation. Each exposure day consisted of placing the animals in ethanol vapor chambers for 14 hrs followed by 10 hrs out of the chambers during which time the animals experience ethanol withdrawal. Control animals underwent similar transport and handling procedures but were exposed to room air instead of ethanol vapor (AIR-exposed). Following 2 weeks of CIE exposure, animals proceeded to the withdrawal drinking phase. During this phase, 3 days of CIE exposure were replaced with 3 days of 2BC withdrawal drinking, which consisted of 14 hr access to 20% ethanol and water in the homecage beginning 10 hrs into withdrawal following removal from vapor chambers. Each withdrawal drinking day was preceded and followed by a day of CIE exposure. Chemogenetic inhibition test sessions began after the 1^st^ week of withdrawal drinking to allow CIE-exposed rats to first achieve stable ethanol intake levels. DCZ, CNO, or vehicle injections occurred 30-45 minutes prior to the onset of withdrawal drinking and ethanol intake was measured 2 hrs after bottle onset. Test sessions were interleaved with at least one non-treatment drinking session to prevent any cumulative effects of repeated drug injections on ethanol intake. Tail-vein blood samples were collected 2 hrs after a DCZ-treated and non-drug treated session for determination of blood ethanol concentrations (BECs) using an Analox alcohol analyzer (Analox Instruments, Atlanta, GA).

### Place conditioning and locomotor activity tests

A two-compartment place preference chamber (Med Associates, ENV-031C) was used to assess whether DCZ-elicited a conditioned place preference or aversion in control rats that had not undergone DREADD stereotaxic surgeries. Rats underwent a 30-minute baseline preference test to assess any innate preference for either compartment. If a rat exhibited a baseline preference (> 60%), the preferred compartment was then paired with injections of DCZ. Otherwise, DCZ or vehicle (saline) injections were randomly paired with a specific compartment. Rats received three 30-minute conditioning sessions for each drug-compartment pairing for a total of six consecutive conditionings sessions. Drug conditioning order was counterbalanced across animals and drug treatment alternated each day. One day after the final conditioning session, rats had a 30-minute, drug-free test session to assess preference for the DCZ-paired compartment.

A separate group of rats was used to assess the effects of DCZ on gross locomotor activity (beam breaks) in the same apparatus used for place conditioning. On the first day, rats underwent a 30-minute session to obtain a baseline measure of locomotor activity.

The following 3 days consisted of saline, DCZ (0.1 mg/kg), and CNO (10 mg/kg) injections, respectively, 30-45 minutes prior to placement in the locomotor chambers. Beam breaks were assessed in 1-minute bins across each 30-minute session, in addition to session totals.

### Statistical Analysis

Mixed effects ANOVAs were used to analyze ethanol intake during dependence-escalated drinking, slice electrophysiology data comparing concentrations of DCZ and CNO on action potential firing, locomotor activity across time, total locomotor activity between drug treatment groups, and preference across time. T-tests were used to analyze ethanol intake and BECs between baseline and DCZ drinking sessions in CIE-exposed rats, slice electrophysiology data assessing effects of DCZ or CNO on number of action potentials relative to baseline, and preference during baseline and post-conditioning test sessions in the DCZ place conditioning experiment. Bonferroni corrections were applied to all post-hoc analyses to correct for multiple comparisons following significant main effects or interactions. All data were analyzed using GraphPad Prism 9.0 software. Data are presented as means ± SEM, and effects were considered statistically significant at *p* ≤ 0.05.

## Results

The goal of the present study was to examine the effectiveness of systemic DCZ for DREADD-based chemogenetic manipulations in behavioral and slice electrophysiological applications in rats. The approach involved examination of hM3Dq-mediated cFos induction by DCZ, acute slice electrophysiological to compare the effects of DCZ and CNO on hM4Di-mediated inhibition of evoked action potential firing, hM4Di-mediated inhibition of an amygdala-dependent alcohol drinking behavior by DCZ, and assessment of the effects of DCZ on gross locomotor activity and place conditioning.

### DCZ-MEDIATED CHEMOGENETIC ACTIVATION ELICITS SELECTIVE CFOS INDUCTION

The first set of experiments assessed whether low dose DCZ is sufficient to support chemogenetic induced of cFos expression. Animals received unilateral injections of an excitatory DREADD (hM3Dq) such that the opposite hemisphere could serve as within-subject control for cFos expression. Upon examination, cFos+ cells were observed in the control hemisphere, indicating that DCZ alone did not induce cFos in cells lacking hM3Dq. Therefore, one sample t-tests were used to compare mean cell counts in the hM3Dq-expressing hemisphere to a theoretical mean of zero. The number of hM3Dq.mCherry+ cells in the z-stack image series was significantly greater than zero (103.0±23.18, t_3_ = 4.443, *p* = 0.021; **Figure 1A, D**), indicating sufficient virus expression in the injected hemisphere. In addition, a one sample t-test revealed the number of cFos+ cells in the hM3Dq-expressing hemisphere were significantly greater than zero (88.50±21.57, t_3_ = 4.102, *p* = 0.026; **Figure 1B, E**). Moreover, the percentage of hM3Dq.mCherry+ cells that were also cFos+ was significantly greater than zero (77.58±3.39, t_3_ = 22.89, *p* = 0.0002; **Figure 1C, F**). Overall, these results suggest that DCZ induces robust cFos expression in cells containing excitatory DREADD receptors without non-specific cFos induction.

**Figure 1.**
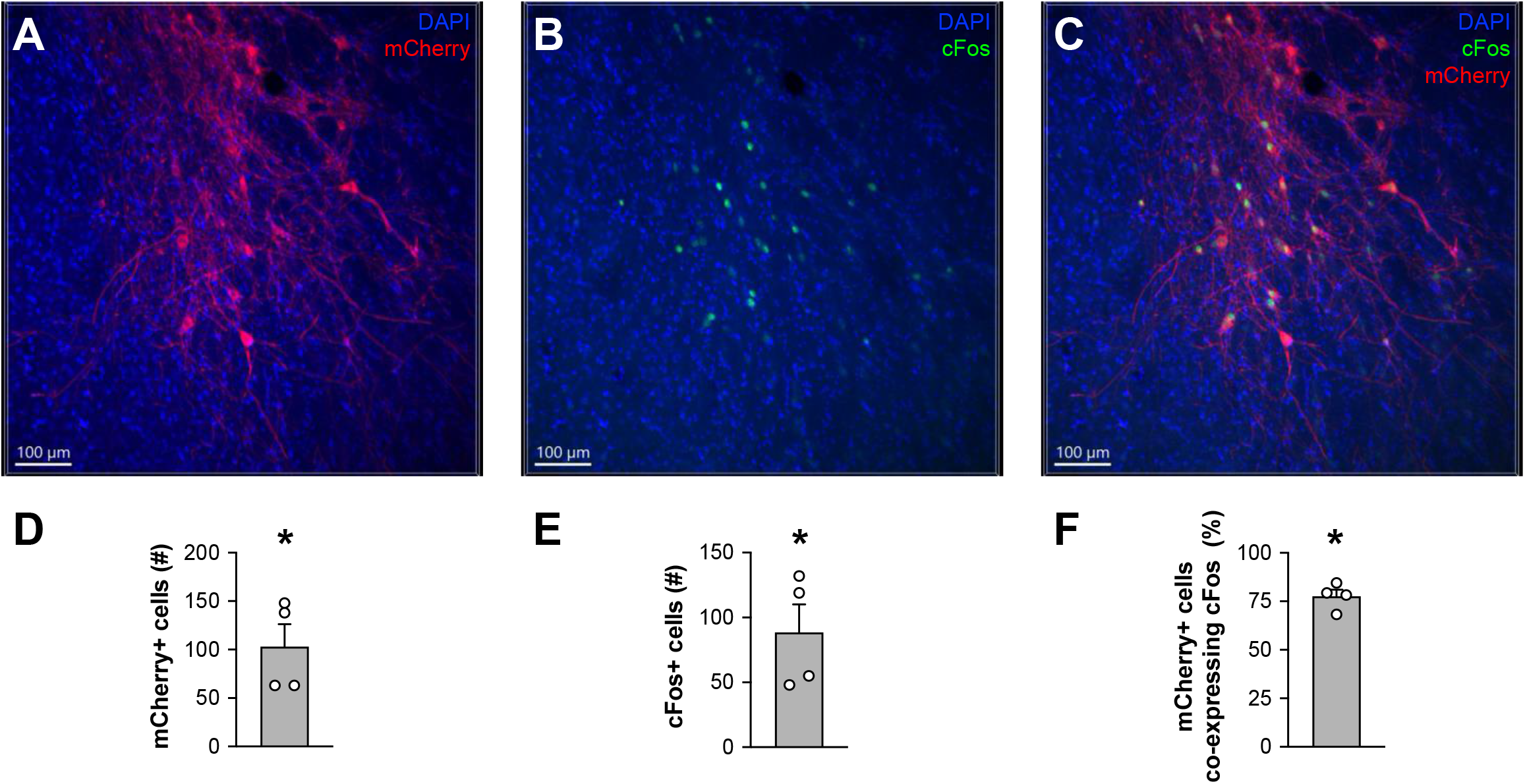
DCZ-mediated chemogenetic activation elicits selective cFos induction in hM3Dq-expressing Cells. Representative 10x images of the CeA containing mCherry+ **(A)**, cFos+ **(B)**, or mCherry+/cFos+ cells **(C)** following injections of 0.1 mg/kg DCZ in male Long-Evans rats with unilateral expression of hSyn.hM3Dq.mCherry. The quantification of average mCherry+ **(D)**, cFos+ **(E)**, and mCherry+/cFos+ **(F)** cell counts are depicted below each image. The opposite hemisphere, that did not receive the hM3Dq virus, did not exhibit any quantifiable cFos+ cells. \**p* < 0.05; *n* = 4 rats.

### CHARACTERISTICS OF DCZ IN SLICE ELECTROPHYSIOLOGY PREPARATIONS

The next set of experiments compared the potency of DCZ with the commonly used DREADD ligand, CNO, for chemogenetic inhibition of neural activity in an acute slice electrophysiology preparation. Neurons in the central amygdala (CeA) expressing the hM4Di underwent whole-cell patch clamp recordings (**Figure 2A**) to determine potency of DCZ and CNO on inhibiting firing rate (**Figure 2B**). A mixed-effects ANOVA with concentration and drug as factors revealed significant main effects of concentration (F_(2.392, 19.14)_ = 22.88, *p* < 0.0001) and drug (F_(1,8)_ = 9.386, *p* = 0.0155), indicating that increasing concentrations of drug further inhibited firing rate and that DCZ produced greater inhibition of firing rate relative to CNO. In addition, there was also a significant drug x concentration interaction (F_(4, 32)_ = 3.191, *p* = 0.0259). Post-hoc analysis revealed that DCZ produced significant inhibition of firing rate at 500 and 1000 nM concentrations, relative to baseline, whereas CNO significantly suppressed firing only at the 1000 nM concentration (all *p* values < 0.05), which suggests that DCZ was more potent than CNO at reducing firing rate in hM4Di+ neurons. We separately plotted action potential firing during baseline and maximal drug concentration (1000 nM) conditions for both DCZ (**Figure 2C**) and CNO (**Figure 2D**). Paired t-tests indicated that both DCZ (t_4_ = 6.318, *p* = 0.0032) and CNO (t_3_ = 4.382, *p* = 0.0220) significantly reduced action potential firing at the 1000 nM concentration.

**Figure 2.**
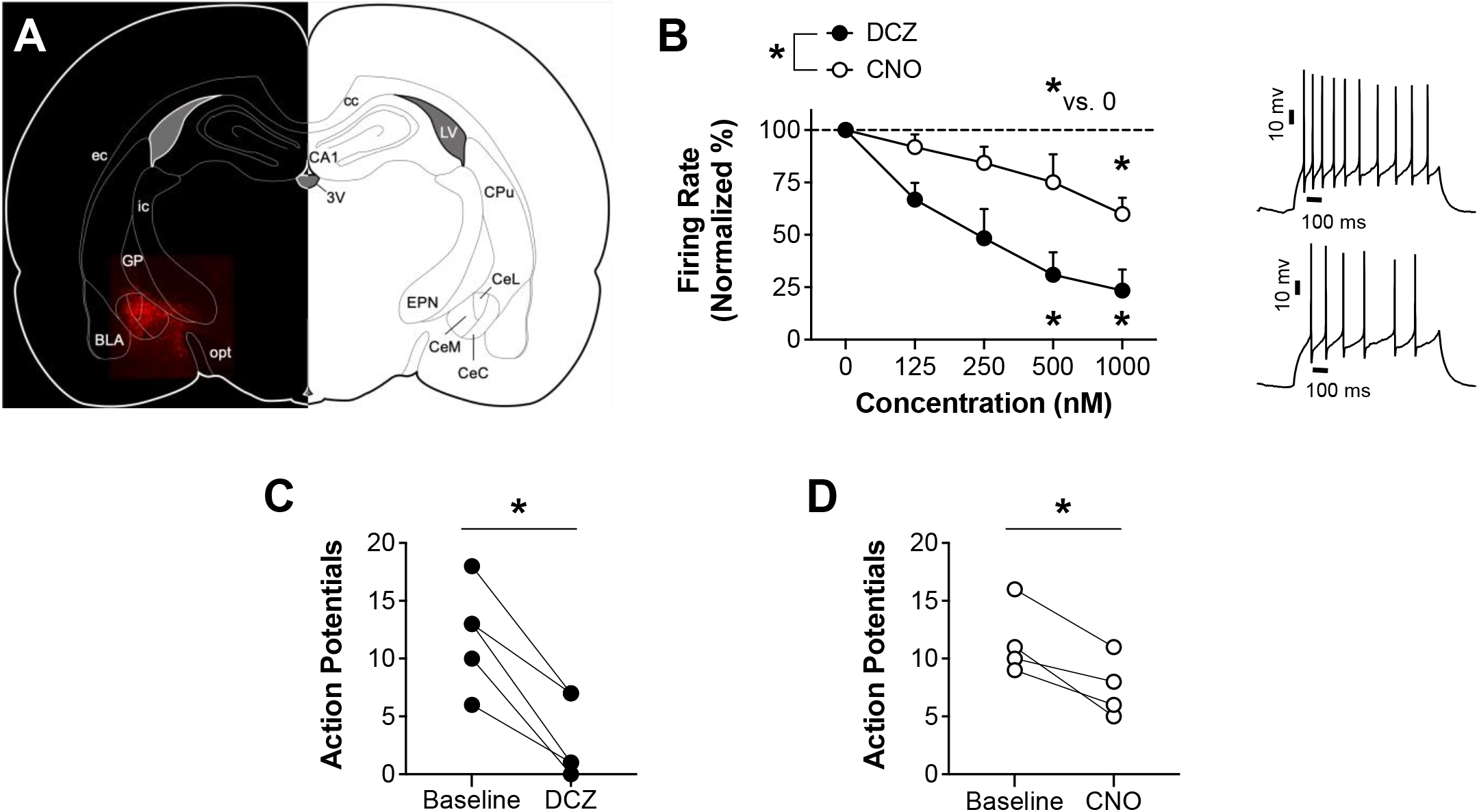
*In vitro* electrophysiology profile of DCZ and CNO in CeA neurons expressing hM4Di DREADDs. **(A)** Atlas schematic depicting viral expression of hM4Di.mCherry in CeA neurons targeted for slice electrophysiology recordings. **(B)** Left: Concentration-response curve depicting percent inhibition of firing rate in CeA neurons expressing hSyn.hM4Di.mCherry during bath application of varying concentrations of either DCZ or CNO. Right: Representative traces of action potential firing in hM4Di-expressing neurons at baseline (top) or during bath application of 125 nM DCZ (bottom). Average changes in action potential firing during 1 uM application of DCZ **(C)** or CNO **(D)** in individual animals. \**p* < 0.05; *n* = 4-5 rats.

### DCZ-MEDIATED CHEMOGENETIC INHIBITION OF THE CENTRAL AMYGDALA REDUCES DEPENDENT ETHANOL DRINKING

A large number of studies have previously demonstrated that the CeA is necessary for elevated ethanol intake in ethanol dependent animals. Therefore, the next set of studies leveraged this CeA-dependent behavior to assess the efficacy of DCZ for chemogenetic inhibition to reduce ethanol intake. As shown in **Figure 3A**, our behavioral model of ethanol dependence produces robust escalation of ethanol intake in dependent rats. A mixed-effects ANOVA with drinking session and exposure condition as factors revealed significant main effects of session (F_(2.248,21.73)_ = 4.330, *p* = 0.0227) and condition (F_(1, 10)_ = 9.759, *p* =0.0108), and a significant session by condition interaction (F_(3,29)_ = 4.319, *p* = 0.0124). Post-hoc analysis revealed that CIE-exposed rats exhibited significantly escalated ethanol intake starting in sessions 2 and 3, relative to their baseline intake (all *p* values < 0.05), whereas intake by AIR-exposed rats remained unchanged (all *p* values < 0.05). Following escalation of ethanol drinking, we tested the effects of chemogenetic inhibition via DCZ or CNO on ethanol intake (**Figure 3B**). A mixed-effects ANOVA with drug and exposure condition as factors revealed main effects of drug treatment (F_(2.241,19.42)_ = 16.87, *p* < 0.0001) and exposure condition (F_(1,10)_ = 34.43, *p* = 0.0002). Post-hoc analysis revealed that DCZ and CNO reduced ethanol intake in dependent and non-dependent rats to a similar magnitude, relative to their vehicle controls, saline and DMSO, respectively (all *p* values < 0.05). In a separate session, we also assessed whether the DCZ-mediated reduction in ethanol drinking was associated with a reduction in BECs (**Figure 3C-D**). Paired t-tests revealed that DCZ-treated rats exhibited a significant reduction in BECs (t_7_ = 6.988, *p* = 0.0002) and ethanol intake (t_7_ = 3.629, *p* = 0.0084), replicating the results shown in **Figure 3B** and suggesting that DCZ-mediated chemogenetic inhibition of the CeA produces relevant reductions in ethanol intake. Together, these results demonstrate that a hundred-fold lower dose of DCZ is as effective as CNO in supporting chemogenetic inhibition of a CeA-dependent behavior.

**Figure 3.**
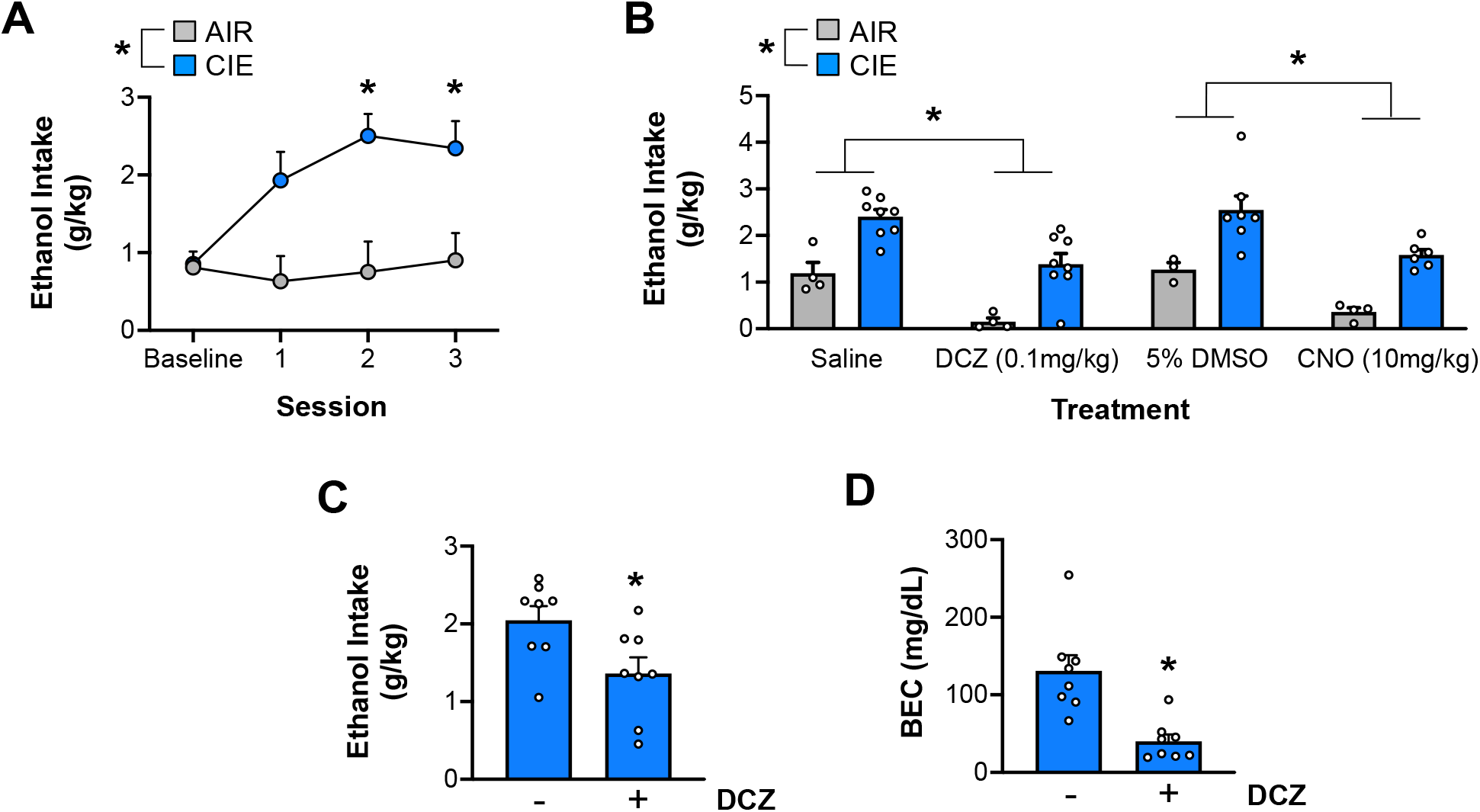
DCZ-mediated chemogenetic inhibition of the central amygdala reduces dependence ethanol drinking. **(A)** Average 2 hr ethanol intake in AIR- and CIE-exposed rats during baseline drinking and 3 sessions of 2BC drinking in the withdrawal drinking phase following CIE exposure. CIE-exposed rats exhibit significantly escalated ethanol intake beginning in session 2, relative to their baseline intake and AIR controls. **(B)** Average 2 hr ethanol intake in AIR- and CIE-exposed rats during drug treatment test sessions. CIE-exposed rats continued to exhibit elevated ethanol intake, relative to AIR controls as suggested by a significant main effect of exposure condition. Further, post hoc analysis following a significant main effect of drug treatment revealed that DCZ (0.1 mg/kg) and CNO (10 mg/kg) significantly reduced ethanol intake in both AIR- and CIE-exposed rats, relative to their vehicle conditions, saline and 5% DMSO, respectively. Moreover, there was no significant difference between ethanol intake in DCZ and CNO sessions. Tail blood was collected from additional non-drug and DCZ (0.1 mg/kg) withdrawal drinking sessions to determine whether the DCZ-mediated reduction in ethanol intake **(C)** also resulted in a concomitant reduction in BECs **(D)**. We replicated the significant reduction in ethanol intake via DCZ-mediated DREADD inhibition **(C)**, which is associated with a significant reduction in blood ethanol levels following 2 hr ethanol access **(D)**. \**p* < 0.05; *n* = 12 rats (AIR = 4, CIE = 8).

### DCZ ALONE DOES NOT ALTER GROSS LOCOMOTOR ACTIVITY OR INDUCE PLACE PREFERENCE/AVERSION

The next set of studies examined whether systemic administration of DCZ alone is associated with non-specific behavioral effects that have been observed with CNO. Toward that end, we assessed the effects of DCZ on locomotor activity (**Figure 4A-B**) and place conditioning (**Figure 4C-D**) in naïve rats not expressing DREADDs. A mixed-effects ANOVA of locomotor activity with time and drug treatment as factors revealed a significant main effect of time (F_(6.664,79.96)_ = 5.479, *p* < 0.0001), but no significant main effect of drug treatment (F_(3,12)_ = 1.100, *p* = 0.3871) or drug treatment x time interaction (F_(87,348)_ = 1.191, *p* = 0.1399). These results reveal that locomotor activity decreased across the 30-minute session in all groups, regardless of drug treatment, suggesting that DCZ does not alter gross locomotor activity. A mixed-effects ANOVA of percent preference for the DCZ-paired side with time and session (Baseline vs Post-conditioning test) as factors yielded no significant main effect of time (F_(2.857, 28.57)_ = 2.561, *p* = 0.0771), session (F_(1,10)_ = 0.07812, *p* = 0.7856), or interaction (F_(5,50)_ = 1.681, *p* = 0.1564), indicating that multiple injections of DCZ in animals without DREADDs is neither rewarding or aversive.

**Figure 4.**
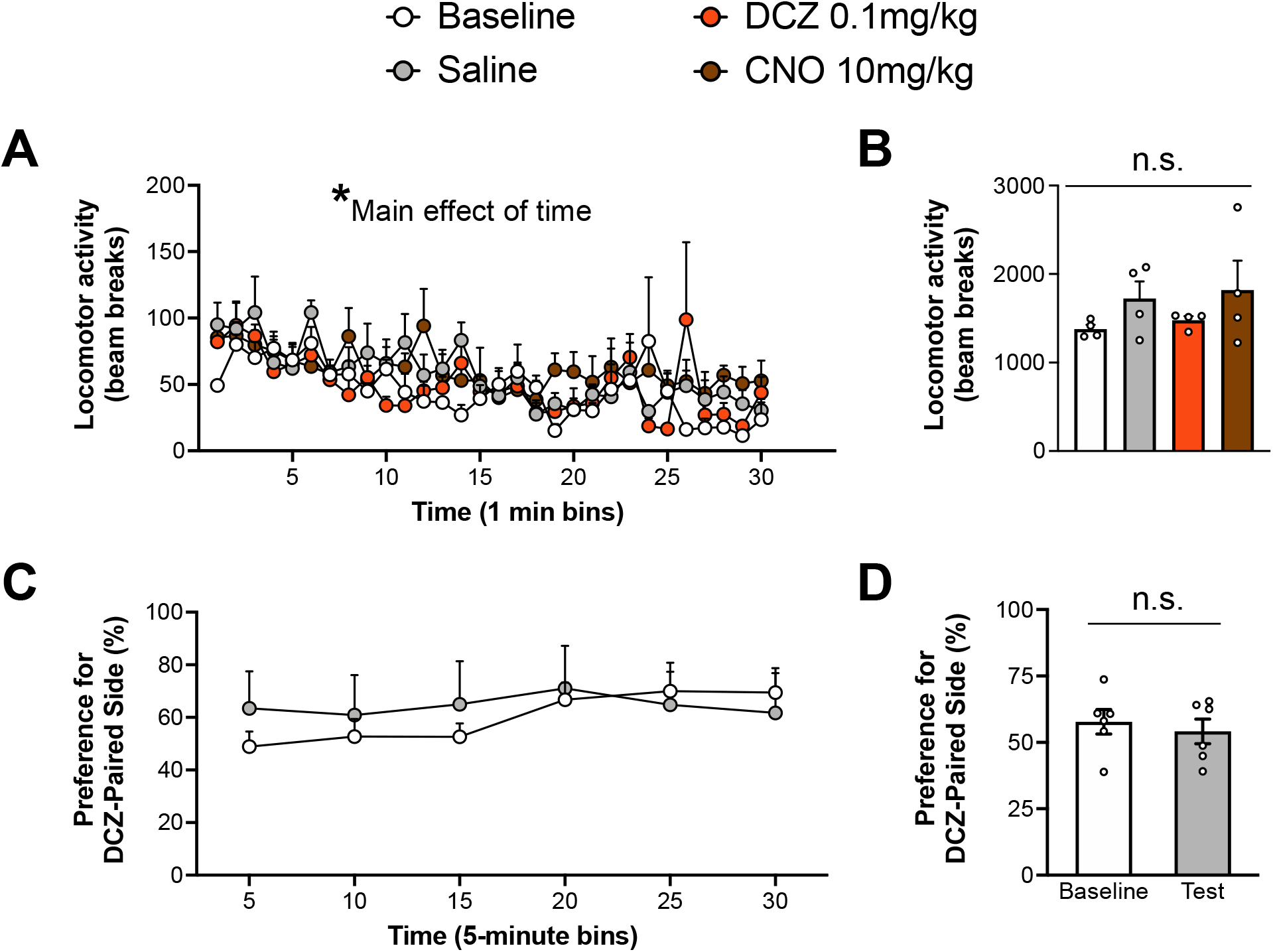
DCZ does not alter gross locomotor behavior and does not condition a place preference/aversion in naïve rats without DREADD expression. **(A)** Mean locomotor activity across 30-minute sessions during baseline or drug treatment conditions. **(B)** Mean total locomotor activity for baseline and drug treatment conditions. There was no significant difference in total locomotor activity between conditions **(C)** Mean percent preference for the DCZ-paired side across 30-minute baseline and post-conditioning test sessions. **(D)** Mean total preference for the DCZ-paired side during baseline and the post-conditioning test. There was no significant difference in time spent in the DCZ-paired side between baseline and post-conditioning test sessions. \**p* < 0.05; *n* = 4 (locomotor activity) and 6 (place conditioning) rats.

## Discussion

The present study provides the first evidence of behavioral and *in vitro* slice electrophysiological efficacy of the DREADD ligand, DCZ, in rats. Our data show that a low dose of DCZ (0.1 mg/kg) administered systemcially effectively supports hM3Dq and hM4Di DREADD-mediated behavior and electrophysiological effects without apparent confounding, non-specific alterations in locomotor behavior or associatve conditioning. Systemic administration of DCZ to selectively induced robust cFos expression in hM3Dq-expressing neurons without producing any detectable cFos in the uninfected control hemisphere. In an *in vitro* slice electrophysiological preparation, DCZ effectively inhibited neuronal firing in hM4Di-expressing neurons. Moreover, DCZ produced greater inhibition of action potential firing at a lower concentration, compared to CNO. Systemic DCZ-mediated hM4Di inhibition reduced ethanol intake in ethanol dependent and non-dependent rats expressing hM4Di in the central amygdala. Lastly, systemic DCZ did not alter gross locomotor activity or produce a conditioned place preference/aversion in control animals that lacked DREADD expression, providing evidence that DCZ alone does not induce noticeable off-target behavioral alterations. Taken together, our findings suggest that DCZ is an effective ligand for use with hM3Dq and hM4Di DREADDs in rats and supports prior evidence that DCZ is a superior DREADD ligand that does not appear to produce off-target behavioral or physiological outcomes.

The goal of the hM3Dq cFos experiment was to assess whether low dose DCZ could produce a robust and selective activation of cells expressing the excitatory DREADD, hM3Dq in rats. We chose cFos immunolabeling as the readout because induction of cFos protein is a widely used index of neuronal activity with a temporally defined expression window (Dragunow & Faull, 1989; Kovács, 2008). DCZ-mediated activation of hM3Dq resulted in robust induction of cFos expression in hM3Dq-expressing cells and surrounding non-infected cells in the viral injected hemisphere, but did not induce any detectable cFos expression in the opposite non-injected control hemisphere. Because identical imaging settings were used for both hemispheres, it is possible that DCZ induced very low levels of cFos in the control hemisphere that we did not detect. However, given the robust staining in the virus hemisphere, any sparse cFos expression in other brain regions is likely negligible. Overall, results from this set of experiments reveal that systemically adminstered DCZ robustly and selectively activates hM3Dq-expressing neurons *in vivo*.

Prior studies that assessed hM4Di-mediated inhibition with CNO in slice electrophysiology applications typically monitored changes in membrane potential, action potential firing, rheobase, etc., and reported near-maximal inhibitory responses with 5 or 10 uM CNO (Anderson et al., 2019; Buchta et al., 2017; H. Li et al., 2013; M. M. Li et al., 2019). The highest CNO concentration used in the present study was 1 uM, which produced ~40% inhibition of firing. This result appears appropriate relative to studies using 5 or 10-fold higher CNO concentrations that appear to achieve near-maximal inhibition. However, the same concentration of DCZ (1 uM) produced ~75% percent inhibition, providing further support that DCZ offers better potency for hM4Di-mediation inhibition, relative to CNO.

For the hM4Di DCZ behavioral experiments, we chose to examine ethanol dependence-induced escalation of drinking as a number of previous studies have consistently demonstrated that the central amygdala is necessary for escalated intake in ethanol dependent animals (de Guglielmo et al., 2019; Hyytiä & Koob, 1995; Vendruscolo & Roberts, 2014). To our knowledge, this is the first report to examine the effect of DREADD hM4Di inhibition of the central amygdala on dependence-induced escalation of drinking. While we observed that both DCZ and CNO-mediated hM4Di inhibition of the central amygdala reduced ethanol intake to a similar extent, DCZ was effective at a 100-fold lower dose. This finding is consistent with the slice electrophysiological observations of concentration differences and supports the contention that DCZ is a superior DREADD ligand (Nagai et al., 2020). In addition to a reduction in ethanol intake in dependent rats in response to systemic administration of either DCZ or CNO, we also observed a significant reduction in ethanol consumption in the non-dependent rats that had not undergone CIE exposure. A prior study reported a similar reduction in ethanol intake in non-dependent rats using a chemogenetic Daun02 inactivation approach to inhibit central amygdala neuronal ensembles (de Guglielmo et al., 2016). Therefore, these observations are consistent with the suggestion that while global inhibition of the CeA may reduce ethanol intake regardless of dependence status, inhibition of specific cell-types and circuits in the CeA may selectively reduce ethanol intake in dependent rats (de Guglielmo et al., 2019).

The present study also assessed whether systemic DCZ used in the prior experiments produced changes in gross locomotor behavior or was intrinsically rewarding/aversive in control rats lacking DREADD expression. Locomotor activity was utilized as it is a high-throughput assay that captures obvious changes in motor function. Furthermore, prior reports have observed changes in gross locomotor activity following administration of other DREADD ligands (Ilg et al., 2018; MacLaren et al., 2016). However, it should be noted that while we did not observe any gross changes in locomotor activity following administration of DCZ, we cannot rule out potential changes to fine locomotor movements that would be captured by more nuanced analyses.

To assess whether DCZ is intrinsically rewarding or aversive we used a place conditioning assay to determine whether repeated injections of DCZ paired with a distinct context could produce a place preference/aversion (Cunningham et al., 2006; McKendrick & Graziane, 2020). The results of these experiments revealed that DCZ did not support a conditioned place preference or aversion of the DCZ-paired compartment. Thus, systemic administration of the same low dose of DCZ that was shown to be an effective agonist of DREADD receptors in the cFos induction experiments, slice electrophysiological analysis of neuronal firing, and ethanol drinking, did not exhibit intrinsic rewarding or aversive properties. These control experiments provide encouraging evidence that DCZ alone does not produce obvious non-specific behavioral effects.

In summary, our findings demonstrate that systemic DCZ is effective for hM3Dq and hM4Di DREADD activation studies in rats. Compared to other available DREADD ligands, DCZ is a promising tool with advantages including increased potency and lack of apparent off-target effects at concentrations that we demonstrate are effective in-vivo for DREADD activation. Moreover, DCZ provides a substantial cost savings benefit due to the low dose requirement relative to CNO and other DREADD ligands. Our observations in rats are consistent with the original findings in primates and mice (Nagai et al., 2020), that DCZ is a clear upgrade from past DREADD ligands and should provide researchers with an improved tool for DREADD activation.

## Acknowledgments

This work was supported by funding from the National Institutes on Alcohol Abuse and Alcoholism grants AA019967 and AA027706 (LJC), T32 AA007474 (TBN, JDO), and F31 AA029622 (TBN).

